# Proof of concept for quantitative urine NMR metabolomics pipeline for large-scale epidemiology and genetics

**DOI:** 10.1101/288993

**Authors:** Tuulia Tynkkynen, Qin Wang, Jussi Ekholm, Olga Anufrieva, Pauli Ohukainen, Jouko Vepsäläinen, Minna Männikkö, Sirkka Keinänen-Kiukaanniemi, Michael V. Holmes, Matthew Goodwin, Susan Ring, John C. Chambers, Jaspal Kooner, Marjo-Riitta Järvelin, Johannes Kettunen, Michael Hill, George Davey Smith, Mika Ala-Korpela

## Abstract

**Background:** Quantitative molecular data from urine are rare in epidemiology and genetics. NMR spectroscopy could provide these data in high-throughput, and it has already been applied in epidemiological settings to analyse urine samples. However, quantitative protocols for large-scale applications are not available.

**Methods:** We describe in detail how to prepare urine samples and perform NMR experiments to obtain quantitative metabolic information. Semi-automated quantitative lineshape fitting analyses were set up for 43 metabolites and applied to data from various analytical test samples and from 1,004 individuals from a population-based epidemiological cohort. Novel analyses on how urine metabolites associate with quantitative serum NMR metabolomics data (61 metabolic measures; n=995) were performed. In addition, confirmatory genome-wide analyses of urine metabolites were conducted (n=578). The fully automated quantitative regression-based spectral analysis is demonstrated for creatinine and glucose (n= 4,548).

**Results:** Intra-assay metabolite variations were mostly <5% indicating high robustness and accuracy of the urine NMR spectroscopy methodology per se. Intra-individual metabolite variations were large, ranging from 6% to 194%. However, population-based inter-individual metabolite variations were even larger (from 14% to 1655%), providing a sound base for epidemiological applications. Metabolic associations between urine and serum were found clearly weaker than those within serum and within urine, indicating that urinary metabolomics data provide independent metabolic information. Two previous genome-wide hits for formate and 2-hydroxyisobutyrate were replicated at genome-wide significance.

**Conclusions:** Quantitative urine metabolomics data suggest broad novelty for systems epidemiology. A roadmap for an open access methodology is provided.

## Introduction

Metabolomics provides a snapshot of an individual’s physiological state, influenced by genetic and lifestyle factors. Urine is produced from blood by the kidneys and contains both endogenous and exogenous compounds.^1^ Among the biofluids commonly used in epidemiology, urine has several advantages: it is abundant, sterile, and easy to collect.^2^ Urine reflects the function of kidneys, including multiple metabolites from several key biochemical pathways in relation to (patho)physiology and cardiometabolic conditions, gut microbial metabolic activities and short-term food consumption.^1,3-5^ Urine samples therefore contain abundant and underutilized information for epidemiology and for potential translational applications.^6^

NMR spectroscopy provides a comprehensive quantitative approach for urine analysis^1,2,5^ and has the potential to offer fully automated high-throughput experimentation in a cost-effective manner, which would be essential for large-scale systems epidemiology.^7-9^ NMR spectroscopy is highly reproducible and requires only minimal sample preparation. Bouatra et al.^1^ have concluded that NMR may currently be the most comprehensive and certainly the most quantitative approach for urine characterisation. However, the signal assignments and quantifications from urine spectra are complicated by signal overlap, as well as considerable variations in signal positions between spectra due to differences in the chemical properties of the samples, such as pH, ionic strength and concentration of multivalent cations.^2^ Some software applications exist which have been used in the analyses of urine NMR data, but currently none of them provide comprehensive automated quantification of the metabolic information.^10-13^

We introduce here a detailed experimental set-up, including all the key attributes of sample preparation and NMR experimentation, for quantitative high-throughput urinary analyses. We also initially demonstrate how fully automated quantitative analyses perform in the case of urine NMR spectra and propose an open access quantitative pipeline of urine NMR metabolomics to facilitate large-scale studies. We present extensive analytical data on intra-assay, intra-individual and inter-individual variation in urinary metabolites. In addition, we detail the characteristics of quantitative urine metabolite data in epidemiology and present novel analyses regarding how the urine metabolites associate with circulating metabolites and lipids. Confirmative genome-wide analyses are also presented. All data domains substantiate the potential usefulness of quantitative molecular data on urine samples in systems epidemiology.

## Materials and methods

### Urine sample preparation

Urine is waste material and thus, in contrast to blood that is strictly buffered, does not entail similar biological regulation. Therefore, there is considerable variation in pH, ionic strength, concentrations of multivalent cations and metabolite composition between samples and individuals. The variations in pH and ionic strength are minimised by the addition of phosphate buffer to the samples. TSP (2,2,3,3-tetradeutero-3-(trimethylsilyl)-propionic acid) is used as a chemical shift as well as an internal concentration reference. The required sample volume is 800 µl. The sample preparation protocol is performed with an automated liquid handler (PerkinElmer JANUS 8-tip Automated Workstation) enabling preparation of approximately 100 urine samples per hour. Detailed instructions for sample preparation are given in **Supplementary Table S1**. From the methodological perspective, any urine sample is appropriate for analysis, i.e., spot urine, overnight or a 24-h collection. The urine samples in this study were stored at −80°C prior to use.

### NMR measurements

NMR data for 4,549 urine samples in the Northern Finland Birth Cohort 1966 (NFBC66; the cohort description is available as **Supplementary Data**) were measured using a 600 MHz Bruker NMR spectrometer, equipped with a cryoprobe (Bruker Prodigy TCI 600 S3 H&F-C/N-D-05 Z) and an automatic cooled SampleJet sample changer. Use of a 600 MHz spectrometer reduces (but does not eliminate) the signal overlap of urine metabolites. Standard water-suppressed measurements are applied. With this hardware set-up NMR data for over 200 urine samples can be automatically collected in 24h. The detailed NMR measurement protocol and parameters are given in **Supplementary Table S2**. Due to day-to-day and person-to-person variation in the volume of urine, that affect the absolute urine metabolite concentrations, it is important to apply a biologically relevant normalisation method. The standard protocol in the field is to normalise to creatinine. We used this approach here but it would be relevant to test this assumption with forthcoming data in extensive epidemiological cohorts by evaluating and comparing multiple methods, e.g., normalisation to the sum of all or selected metabolites in the sample, and potential new methods.^14,15^

Serum samples (n=5,788) from NFBC66 were analysed using a high-throughput quantitative NMR metabolomics platform originating from our team.^7^ This platform provides simultaneous quantification of routine lipids and lipid concentrations of 14 lipoprotein subclasses and major subfractions, and further quantifies abundant fatty acids, amino acids, ketone bodies and gluconeogenesis-related metabolites in absolute concentration units. This serum NMR metabolomics platform has been available since 2009^7^ and it has been used to analyse around 500,000 samples in extensive epidemiological and genetic studies.^8,9^ Details of the experimentation have been described elsewhere^7,8,16^ and the large-scale epidemiological applications have recently been reviewed.^9^ Sixty-one metabolic measures giving a representative overview of the key metabolic pathways were used here.^8,17-19^ Quantitative urine and serum metabolomics data were available for 995 and quantitative urine metabolomics and genome-wide data for 578 individuals.

### Metabolite quantification in urine samples and analytical issues

We have identified over 100 metabolites (**Supplementary Table S3**) and set-up semi-automated lineshape fitting analyses to quantify 43 of these (**Table 1**). In addition to data from multiple analytical test samples, data from 1,004 urine samples from the NFBC66 were quantified. **Table 1** summarises all these data and gives details on the calculations for intra-assay coefficients of metabolite variations in per cent (CV%s), as well as for intra-individual and inter-individual metabolite variation. These semi-automated quantifications rely on the sophisticated lineshape fitting analysis tools developed for high-precision quantitative NMR spectroscopy.^20,21^ **Figure 1** illustrates the characteristics of urine NMR data and the principles of the lineshape fitting analysis. These analyses are, at best, semi-automated and are typically performed separately for multiple spectral regions, i.e., analysing only one or a few metabolites at a time. Thus, these analyses, while being the most robust available, take a considerable amount of time per sample and require manual control of the analysis parameters as well as assessment of fitting results. Thus, when aiming for large-scale epidemiology, regression analysis types of approaches need to be used.^22,23^ However, robust lineshape fitting analyses form the essential base for eventually automating the quantitative metabolite analyses,^22^ i.e., the extensive and detailed data from the lineshape fitting analyses for the 1,004 NFBC66 urine samples will serve as a training set for the automated regression models to be developed.^23^ The automated quantification protocols to be established for the urine analyses will be similar to those we have successfully used in the case of serum NMR metabolomics.^8,23^ We intend to provide an open-access software for the urinary metabolite quantification via a free website. **Supplementary Figure S1** illustrates the building of automated regression models to quantify urinary creatinine and glucose from the NMR spectra. These models are based on the semi-automated lineshape fitting analyses of 999 urine samples; 5 spectra of the 1,004 available were excluded from the automated modelling at this initial stage due to non-optimal shimming and/or baseline features. **Figure 2** shows the final automated regression models for creatinine and glucose and the distribution for urinary glucose concentration in 4,548 people in NFBC66. One spectrum was rejected at this stage by the automated analysis software due to non-optimal shimming.

**Table 1.**
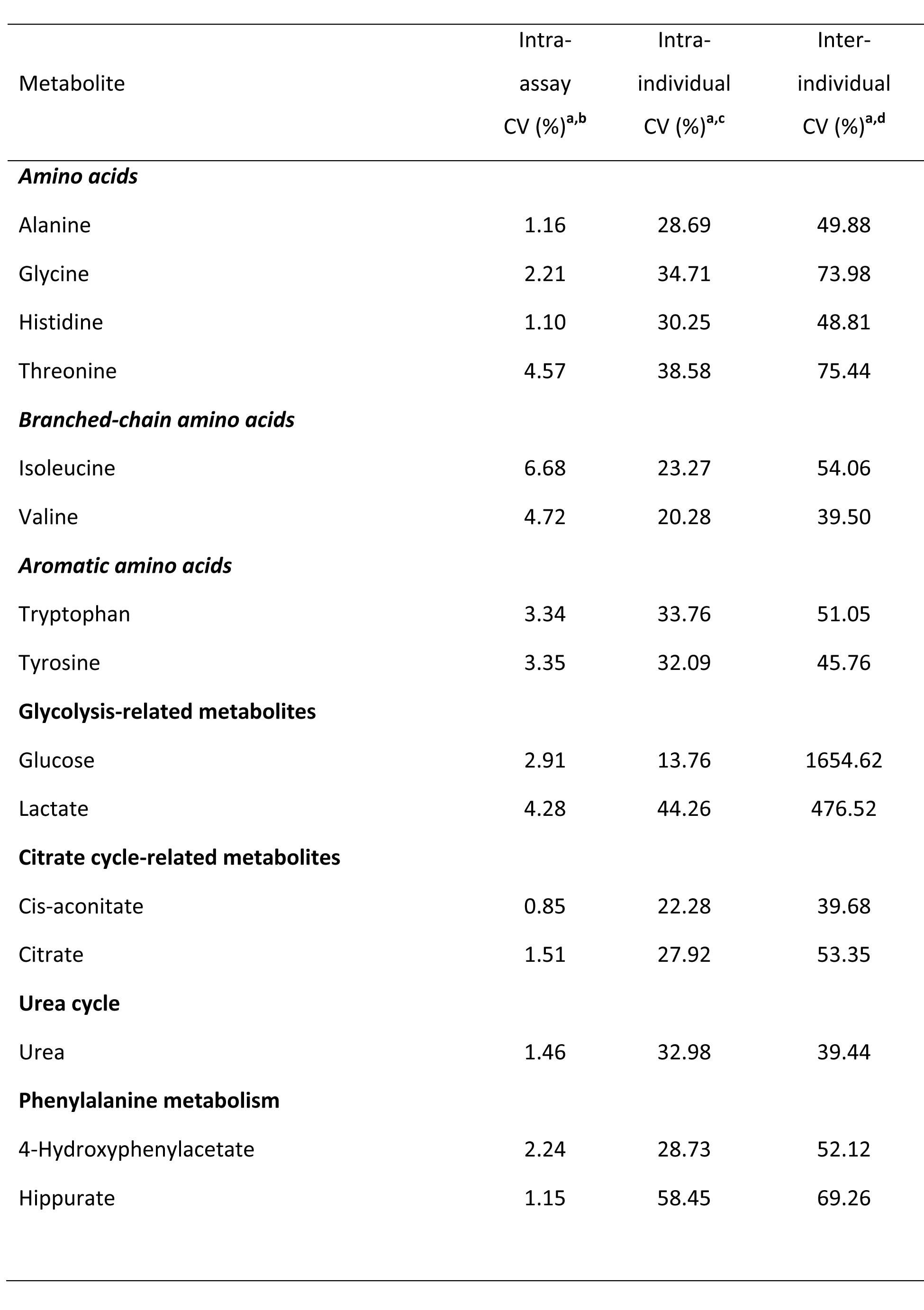
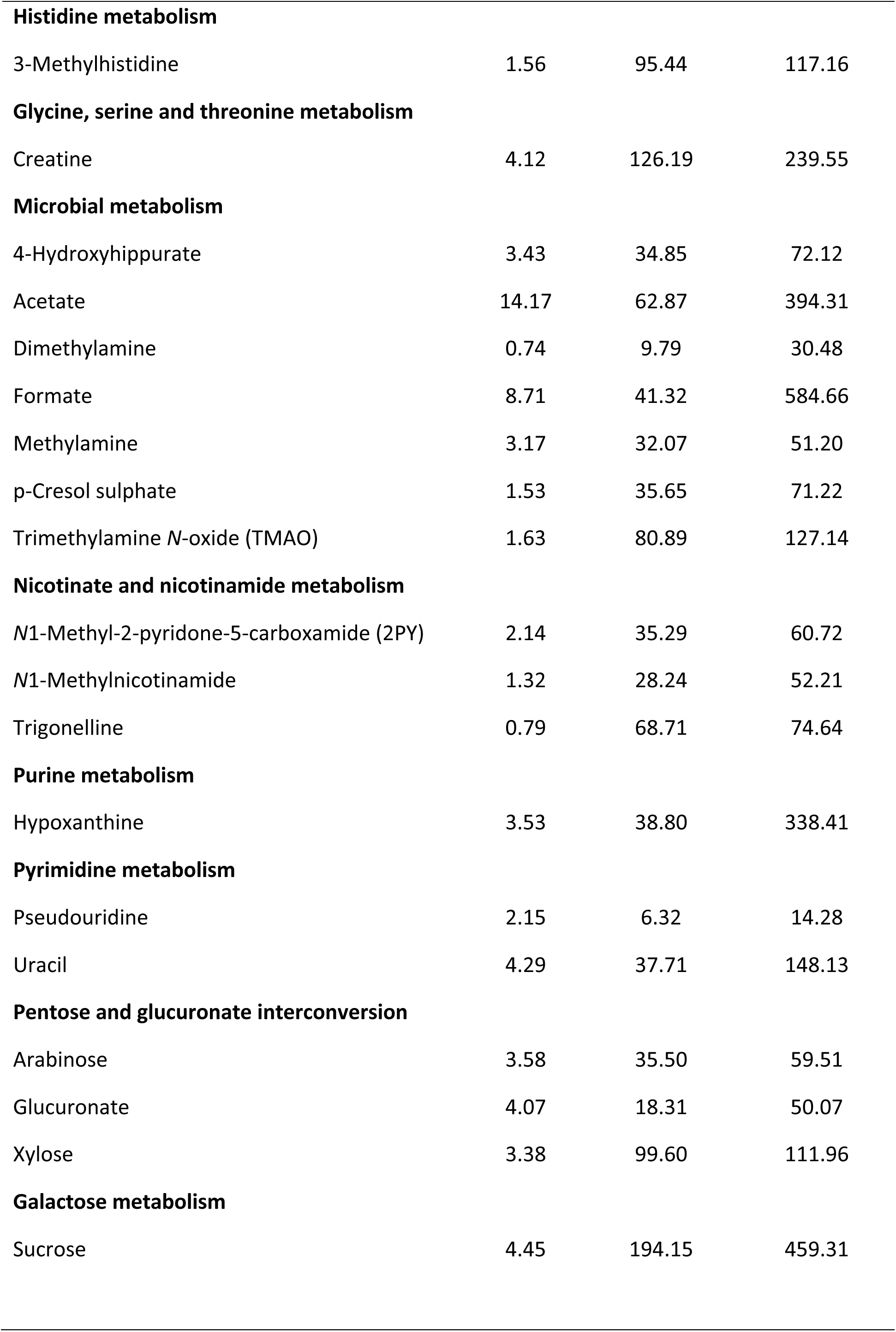
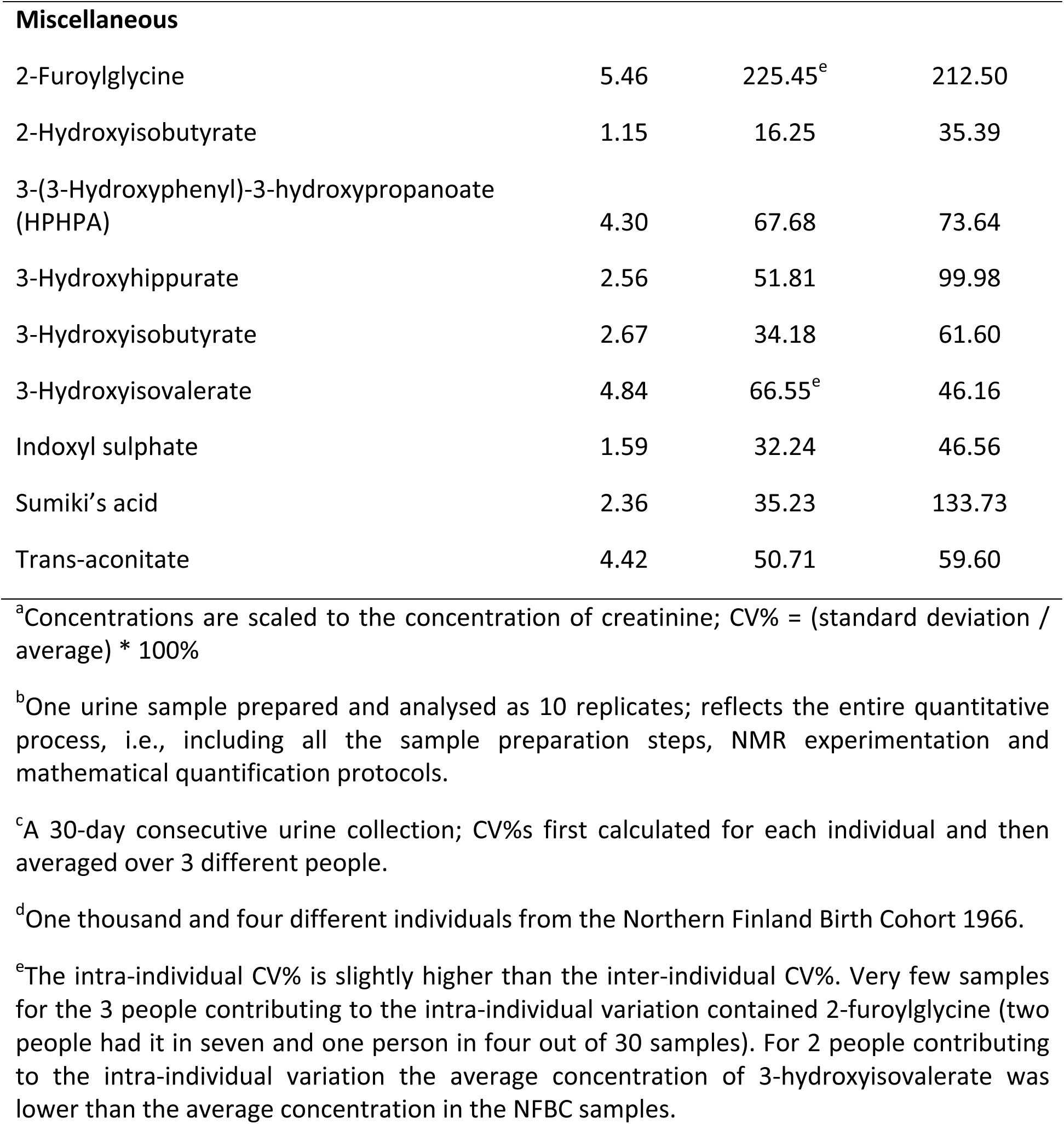
Intra-assay variation as well as intra-individual and inter-individual variation of quantified urine metabolites.

**Figure 1.**
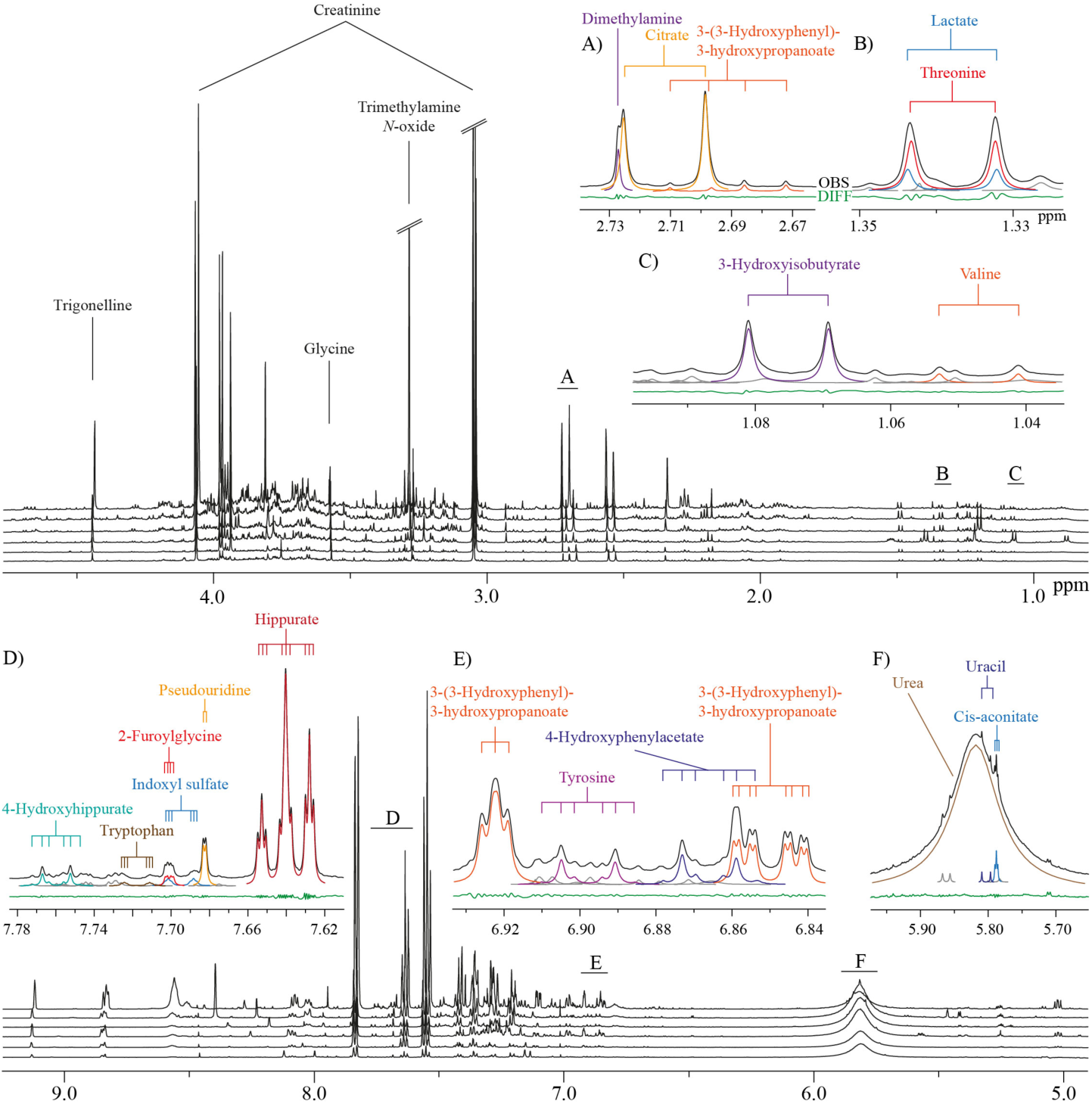
Characteristic ^1^H NMR spectra of human urine from six subjects and illustration of the sophisticated lineshape fitting analyses. Alignment of spectra from six subjects is shown. Heavily overlapping signal structures in multiple areas are typical for these spectra. The insets marked from A to F illustrate how lineshape fitting analyses, incorporating prior knowledge on the individual molecular attributes, can robustly solve the overlap and lead to reliable quantification of the metabolites.^20,21^ Black lines represent the observed spectra and the coloured lines represent the fitted signals. Grey lines indicate currently unidentified signals. The green line at the bottom illustrates the difference between the observed spectrum and the fitted signals. The coupling trees above the spectra demonstrate the multiplet structures directly linked to the molecular attributes and used as constraints in the lineshape fitting analyses.^20,21^

**Figure 2.**
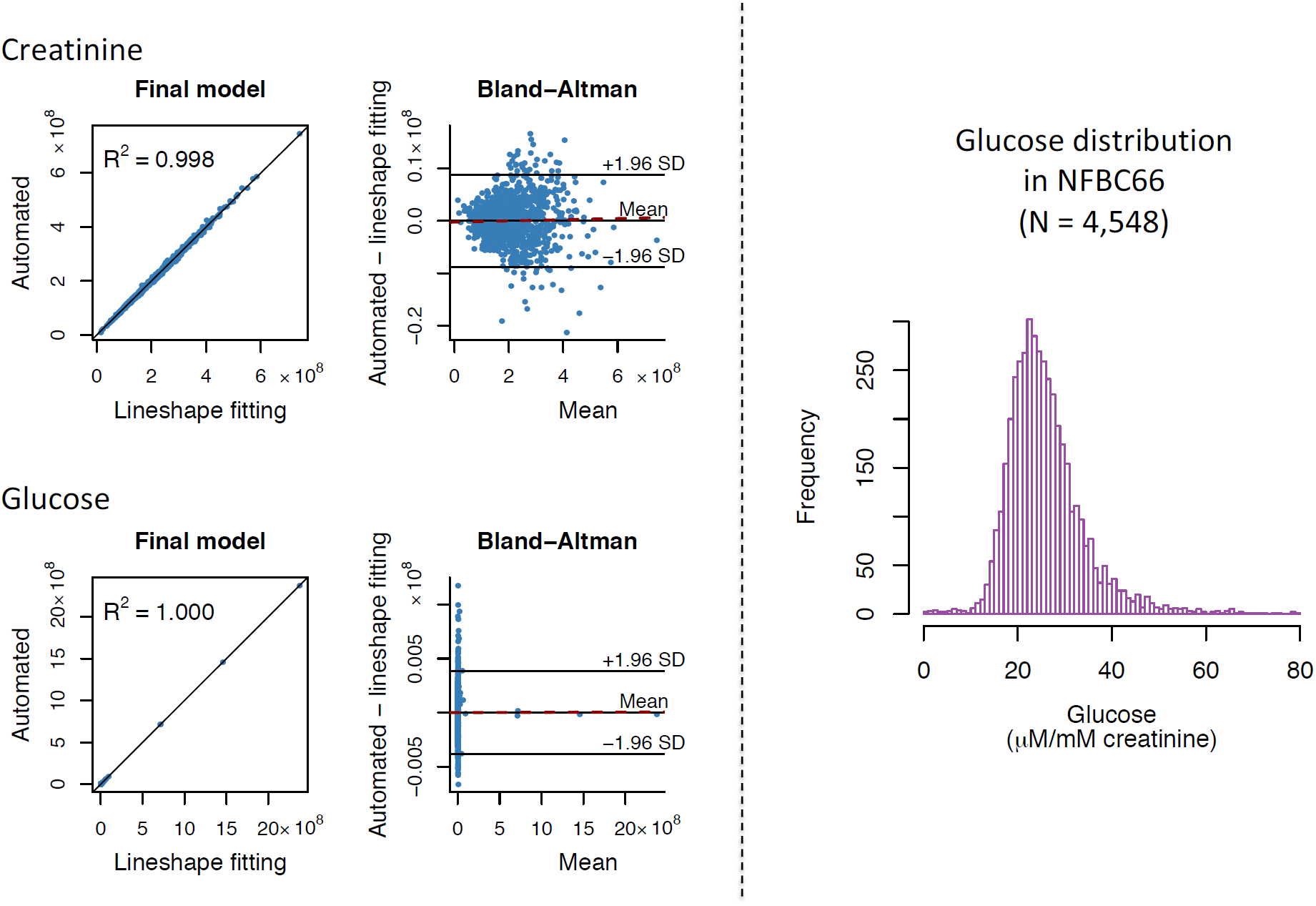
The automated quantification of urinary creatinine and glucose from the NMR spectra. On the left: Building and assessment of the final automated regression models for the absolute signal areas for creatinine and glucose in the NMR spectra (n=999). Training and independent testing results are shown in **Supplementary Figure S1**. In the Bland-Altman plots the solid line in the middle represents the mean bias (between the automated regression and the lineshape fitting analyses results for the absolute signal area) and the two others the mean ± 1.96 SD. The dashed red line represents the regression line for the bias. The equations for the regression lines are *y* = 0.9977*x* + 4.772×10^5^ for creatinine and *y* = 1.000*x* + 31.88 for glucose. Bias as a function of creatinine: *y* = 1.135×10^−3^*x* − 2.390×10^5^ with R^2^ = 0.0006 and bias as a function of glucose: *y* = 2.180×10^−6^*x* − 15.94 with R^2^ = 0.000001. Both automated regression models show excellent quantitative performance and robustness with negligible bias. **On the right:** The distribution of absolute urinary concentration (in µm/mM creatinine) in 4,548 urine samples in NFBC66. The absolute signal areas for the urinary creatinine and glucose used to calculate the distribution are based on fully automated NMR spectral analyses using the final models illustrated on the left. The urinary glucose distribution is positively skewed (88 glucose concentration values > 80 µm/mM creatinine are not drawn for clarity). This is expected due to individuals with prediabetes and diabetes in NFBC66.

### Statistical analyses

Partial correlations adjusted for sex were used to analyse the intra-fluid (urine-urine and serum-serum) and inter-fluid (urine-serum) associations between the quantitative metabolic measures for the NFBC66 samples (N=995). Urine and serum metabolic measures were log-transformed. The results are shown in colour-coded heat maps in **Figure 3** for the intra-serum associations, in **Figure 4** for the intra-urine associations and in **Figure 5** for the inter-fluid urine-serum metabolic associations. The number of principal components (PCs) needed to explain >99% of variation in the metabolic information was 40 PCs for 43 urine metabolites, 27 PCs for 61 serum measures and 66 PCs for the combined metabolic data of 104 metabolic measures. Therefore we used multiple comparison corrected p-value thresholds of 0.001 (i.e., 0.05/40 via the Bonferroni method; P < 0.001 marked with * in the maps), 0.002 (0.05/27, *P < 0.002) and 0.0008 (0.05/66, *P< 0.0008), respectively, to denote evidence in favour of an association.

**Figure 3.**
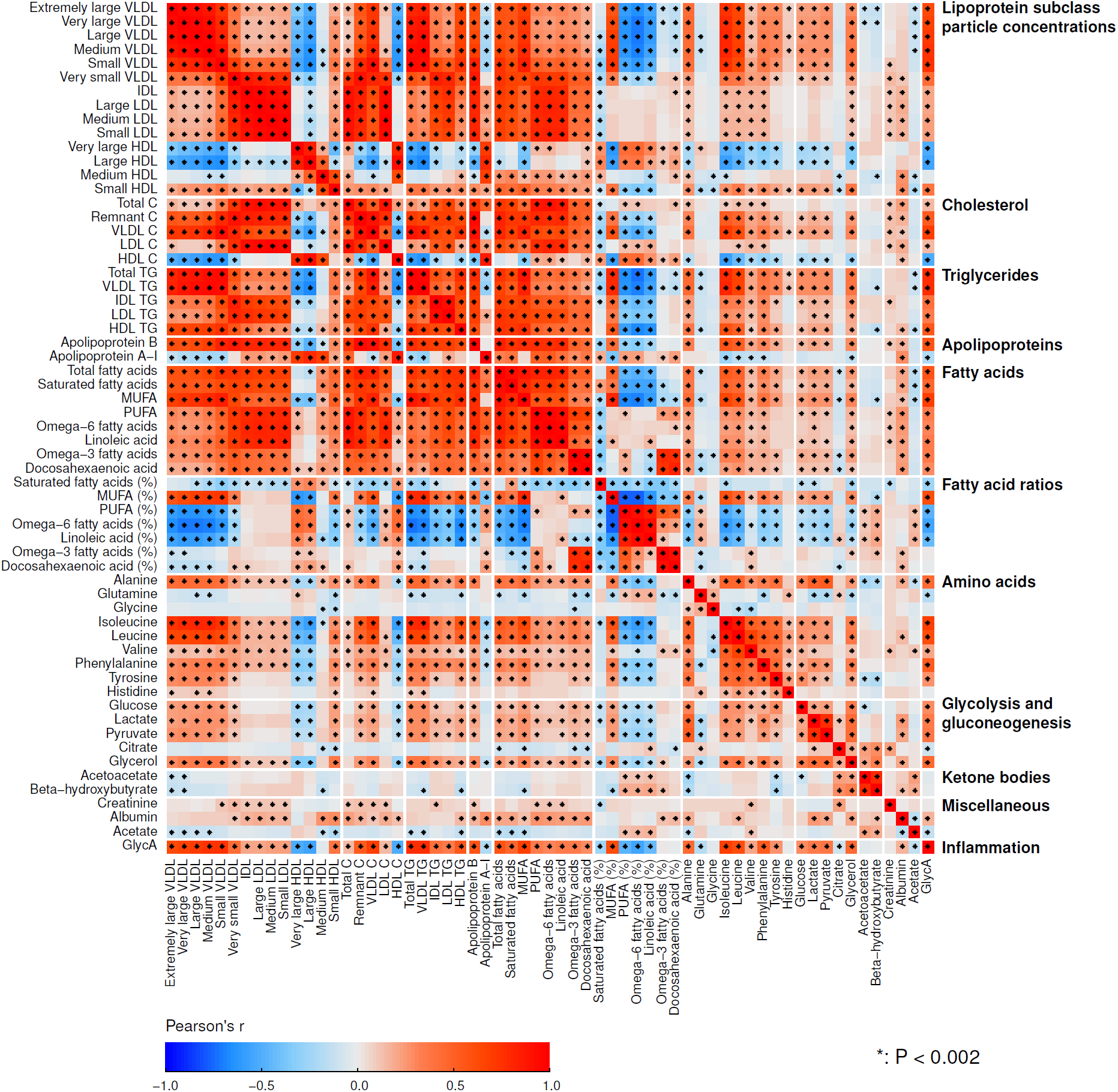
The intra-fluid metabolic associations in serum. The intra-fluid metabolic correlations in serum, i.e., in circulating metabolism, are strong due to multiple key metabolic pathways under heavy systemic control. For example, the metabolism of apoB-containing lipoprotein particles is a continuum and reflected by strong correlations between adjacent lipoprotein subclass particle concentrations. Strong links exist also, e.g., between triglyceride-rich VLDL particles and large cholesterol-rich HDL particles as well as between multiple amino acids.^44^ The colour-coding refers to partial correlations adjusted for sex; n=995 individuals from NFBC66. The heat map is organised manually on the basis of the key metabolic groups and pathways represented by the measures.^17,18^ Twenty-seven principal components were needed to explain >99% of variation in the metabolic information of these 61 serum measures (leading to Bonferroni corrected significance p-value of 0.002 i.e., 0.05/27; marked with * in the map). Abbreviations: VLDL, very-low-density lipoprotein; LDL, low-density lipoprotein; IDL, intermediate-density lipoprotein; HDL, high-density lipoprotein; XXL refers to the largest and XS to the smallest lipoprotein particles in each lipoprotein fraction;^8^ P, particle (concentration); C, cholesterol; TG, triglyceride; PUFA, polyunsaturated fatty acids; MUFA, monounsaturated fatty acids; GlycA, glycoprotein acetyls.

**Figure 4.**
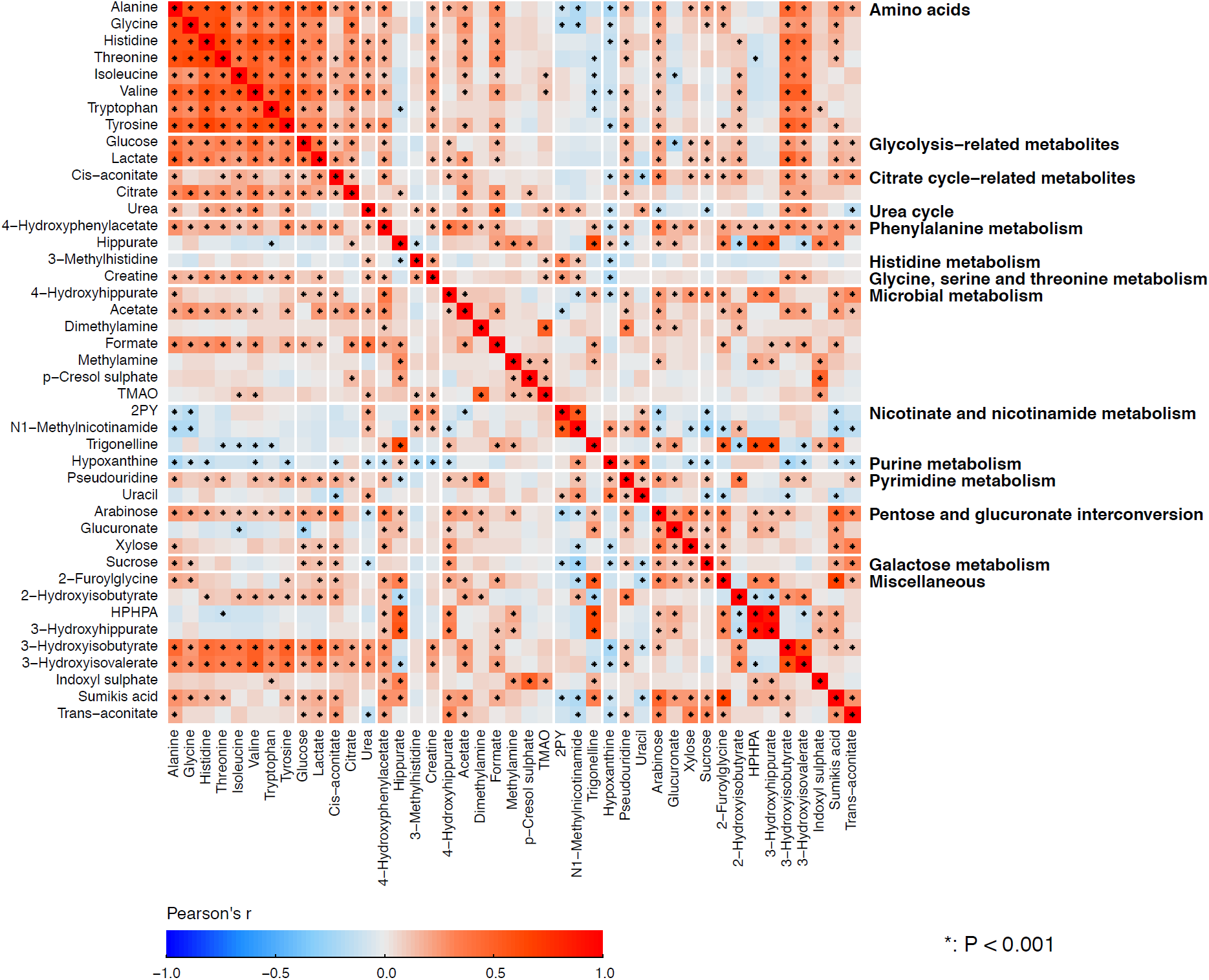
The intra-fluid metabolic associations in urine. The intra-fluid metabolic correlations in urine are generally rather weak and only a few stronger metabolic correlation blocks are noticeable, namely, positive correlations among amino acids, glycolysis- and citrate cycle-related metabolites, 3-hydroxyisobutyrate and 3-hydroxyisovalerate result in clear association clusters. These association characteristics are likely to partly reflect the large intra-individual variation in urinary metabolites, but they are also likely a fundamental sign of metabolic waste with under only limited systemic control. However, the concentrations of the amino acids are rather strongly correlated as would be expected for these apparently healthy individuals with healthy kidneys. The amino acid concentrations also correlate with 3-hydroxyisobutyrate and 3-hydroxyisovalerate, both degradation products of branched-chain amino acids as well as with glucose and lactate, related energy metabolites in gluconeogenesis. Several metabolites related to microbial metabolism are quantified and an interesting correlation cluster is seen between methylamine, p-Cresol sulphate and TMAO. The colour-coding refers to partial correlations adjusted for sex; n=995 individuals from NFBC66. The heat map is organised manually on the basis of the key metabolic groups and pathways represented by the measures (**Table 1**). Forty principal components were needed to explain >99% of variation in the metabolic information of these 43 urine metabolites (leading to Bonferroni corrected significance p-value of 0.001 i.e., 0.05/40; marked with * in the map). Thus, the urine metabolites are generally highly uncorrelated and provide independent metabolic information. Abbreviations: 2PY, *N*1-Methyl-2-pyridone-5-carboxamide; TMAO, Trimethylamine *N*-oxide; HPHPA, 3-(3-Hydroxyphenyl)-3-hydroxypropanoate.

**Figure 5.**
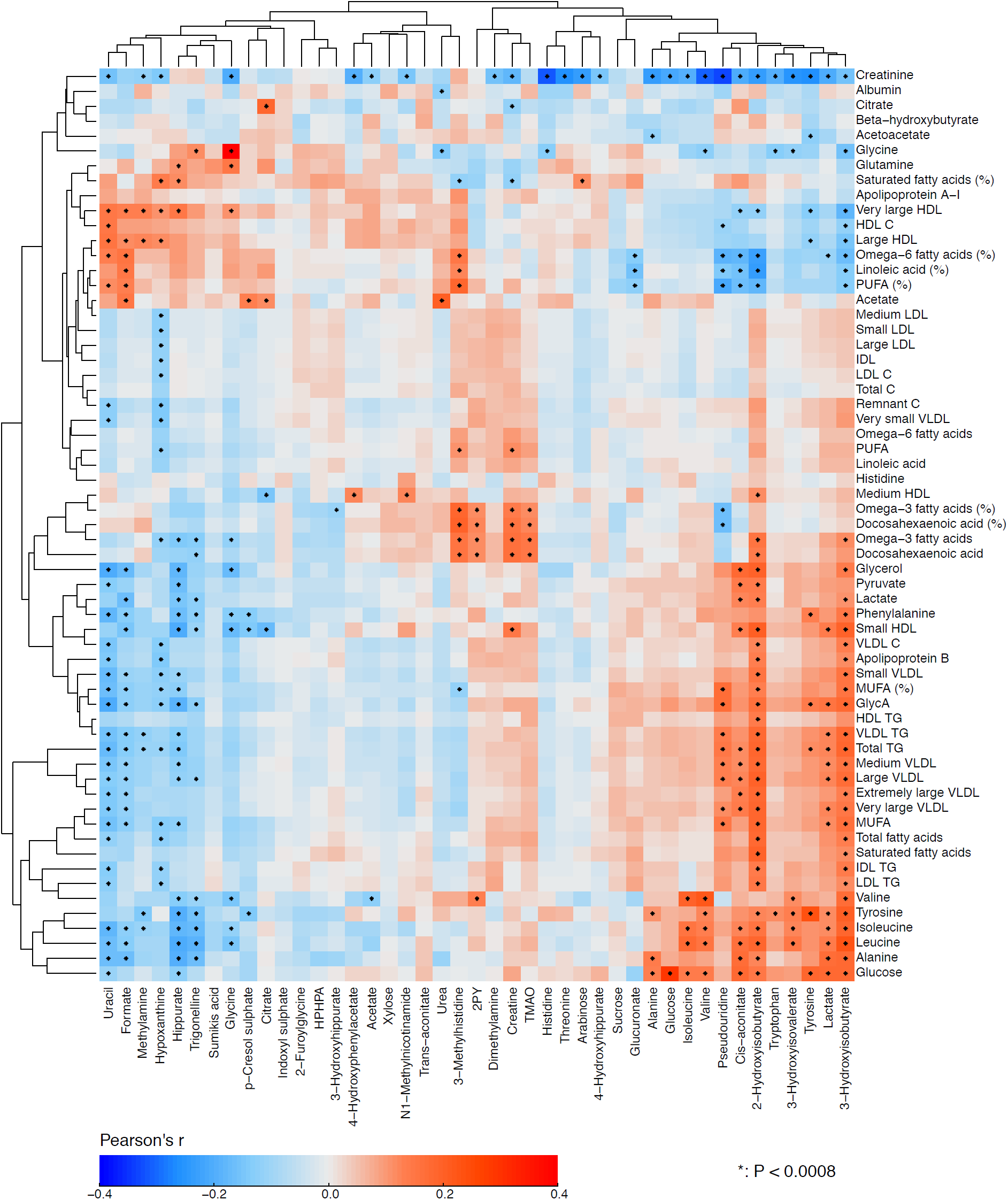
The inter-fluid metabolic associations between urine and serum. The inter-fluid metabolic correlations between urine and serum are rather weak. However, several clearly detectable associations are present. The amino acid concentrations in serum and in urine are strongly positively associated, except for histidine for which the correlation appears very weak. There is an intriguing positive association between urinary TMAO and serum polyunsaturated omega-3 fatty acids. Notably, circulating TMAO has been linked to the pathogenesis of cardiovascular disease.^45^ However, we do not yet have data to associate urinary TMAO concentrations with cardiometabolic outcomes and its concentration in serum is too low to be quantified by serum NMR metabolomics. In addition, serum polyunsaturated omega-6 fatty acids associate negatively with multiple urinary metabolites in relation to amino acid, energy and microbial metabolism, for example, 2-hydroxyisobutyrate, cis-aconitate, and pseudouridine. Multiple urinary metabolites, e.g., 3-hydroxyisobutyrate, lactate, pseudouridine, and cis-aconitate associate with circulating amino acids, glucose and creatinine. For example, for cis-aconitate, a key component in the citric acid cycle, these associations are not unexpected. Cis-aconitate also associates with serum triglycerides. On the other hand, urinary uracil (a naturally occurring pyrimidine found in RNA and, e.g., related to carbohydrate metabolism and sugar transport) is positively associated with serum high-density lipoprotein (HDL) cholesterol. The rationale for this association is not evident; though it could be due to uracil’s involvement in energy metabolism and the inverse association between serum triglycerides and HDL cholesterol. The colour-coding refers to partial correlations adjusted for sex; n=995 individuals from NFBC66. The heat map is organised via 2-dimensional hierarchical clustering. Sixty-six principal components were needed to explain >99% of variation in the metabolic information of these 104 metabolic measures combining the quantitative information from urine and serum (leading to Bonferroni corrected significance p-value of 0.0008 i.e., 0.05/66; marked with * in the map). Combining quantitative urine metabolite data with serum metabolomics would thus evidently increase the independent metabolic information content of the data set. Abbreviations are as detailed in the captions for **Figure 3** and **Figure 4**.

Adiposity is a causal risk factor for many cardiometabolic conditions^24^ and it has been previously studied in relation to urinary metabolites.^4^ Therefore, we wanted a preliminary understanding and comparison of our quantitative urine metabolite data and analysed the associations of BMI with the 43 quantified urine metabolites. A linear regression model was fitted for each outcome measure (concentrations of metabolites in urine and those corresponding in serum) using BMI as the explanatory variable. All metabolic measures were log-transformation and scaled to SD units (by subtracting the mean and dividing by the standard deviation). Association magnitudes are reported in SD units to ease the comparison across multiple measures (**Figure 6**).

**Figure 6.**
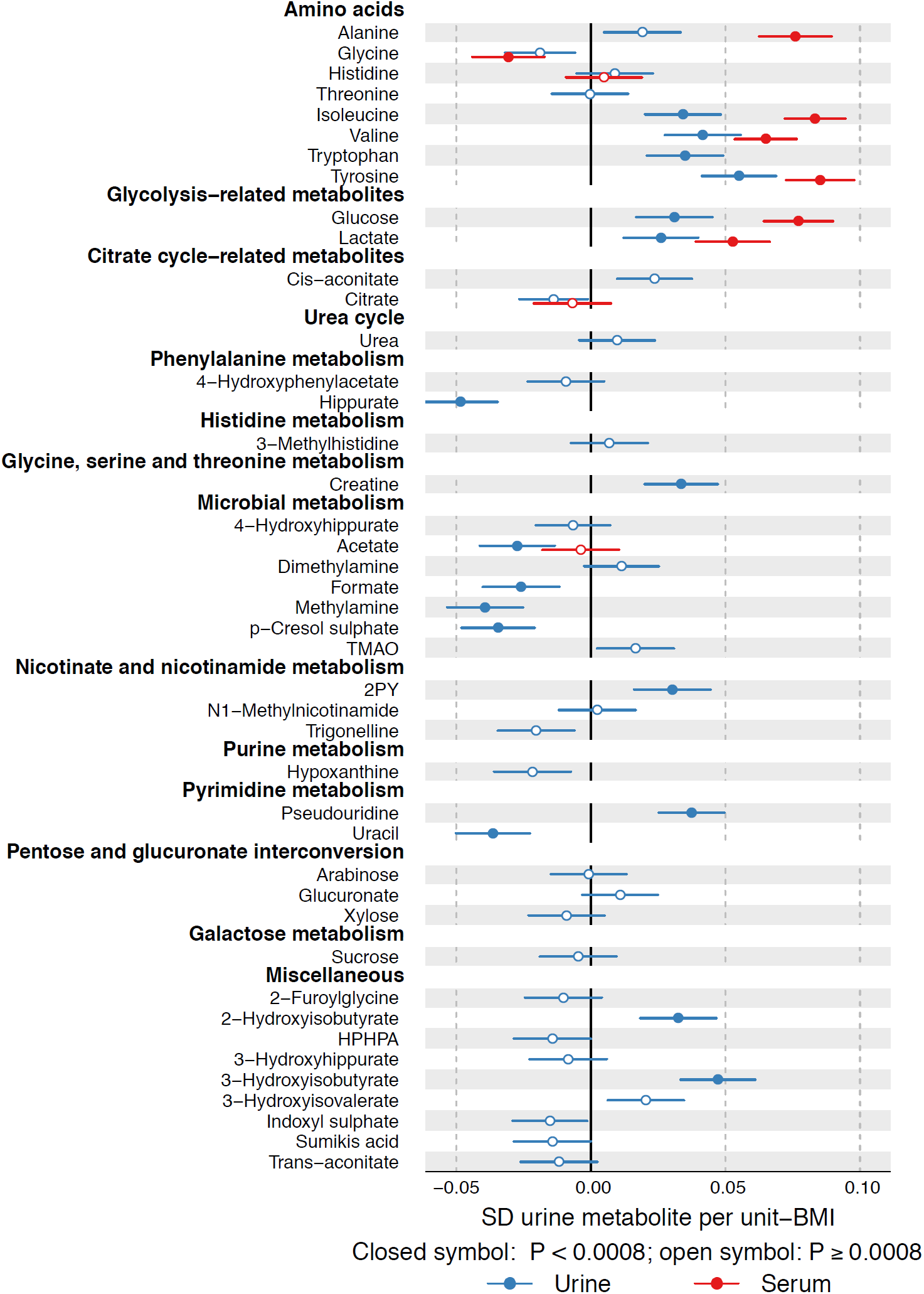
Associations of metabolites quantified in both urine and serum with body mass index. Multiple associations are notable between urinary metabolites and BMI. For example, BMI associates negatively with urinary p-Cresol sulphate and hippurate, and positively with 2-hydroxyisobutyrate and branched-chain amino acids isoleucine and valine, and aromatic amino acids tryptophan and tyrosine. For all the amino acids that are quantified from both urine and serum, the association direction with BMI is the same in serum and in urine; the association strengths however tend to be weaker in urine. Abbreviations are as detailed in the caption for **Figure 4**.

As another positive control for the urine platform, we conducted a genome-wide analysis study (GWAS) of urine metabolites in 578 individuals and compared our results with previous GWA studies. Manhattan plots for formate and 2-hydroxyisobutyrate are shown in **Figure 7** and details of the genetic data and analyses are given in the corresponding caption.

**Figure 7.**
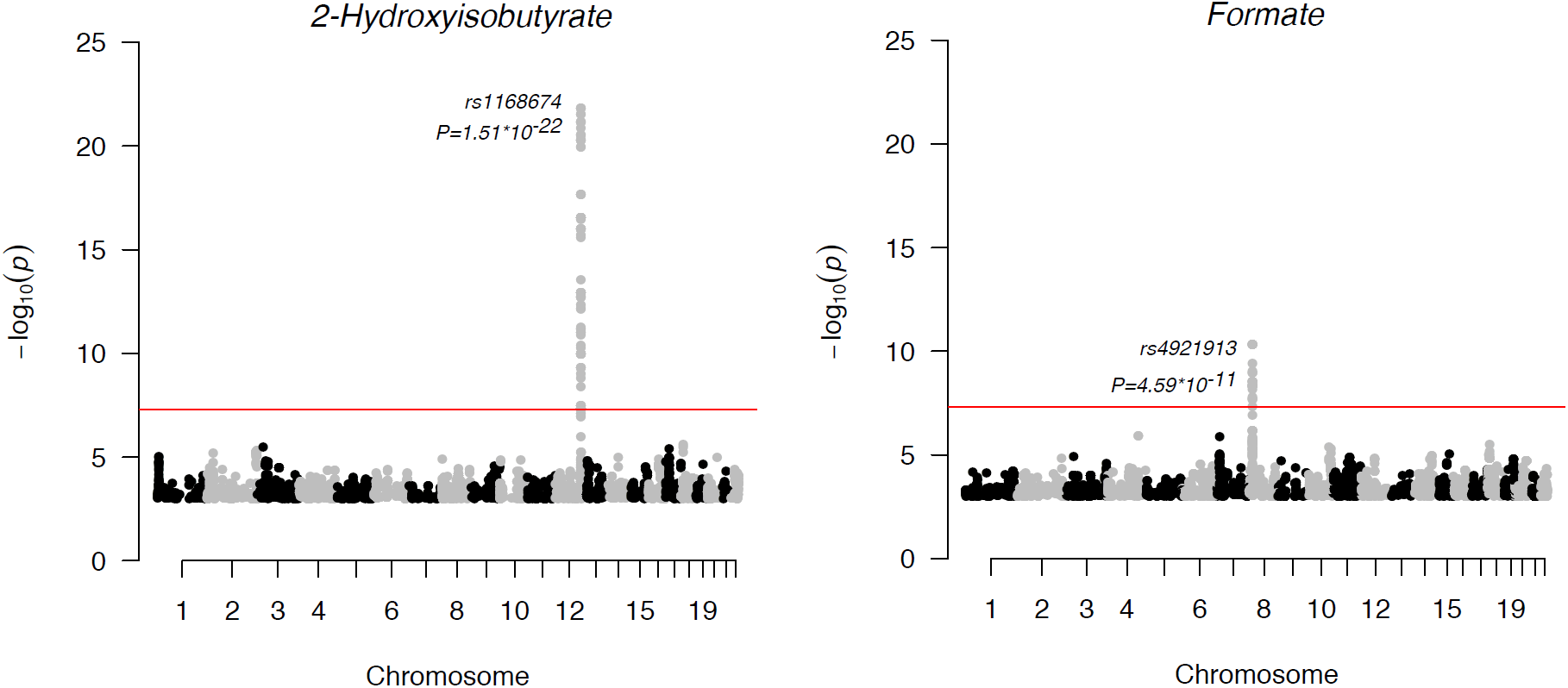
Manhattan plots of the GWAS of formate and 2-Hydroxyisobutyrate. The SNP associations across the whole genome are presented. For plotting purposes, the associations with P-value larger than 1*10^−3^ are not shown. Each dot is a –log_10_ of P-value of the association between the genetic variant and the metabolite using an additive model. The dots are ordered using the chromosome number and base pair position of the variant in the chromosome. Red line indicates the standard level of genome-wide significance (5*10^−8^). The top signals in these two plots were significant after correcting the genome-wide significance threshold for 40 independent tests (P<1.25*10^−9^). All metabolite concentrations were first adjusted for sex, and ten first principal components from genomic data and the resulting residuals were transformed to normal distribution by inverse rank-based normal transformation. NFBC66 was genotyped using Illumina HumanHap 370k array. The genotypes were imputed using the Haplotype Reference Consortium pipeline.^46^ The results were filtered using minor allele frequency cut-off of 5% or greater and imputation info 0.8 or greater. The analysis software was SNPTEST 2.5.1 using additive model for association testing.^47^

## Results and discussion

### Analytical issues in urine metabolomics

**Table 1** lists the currently quantified 43 urine metabolites with their intra-assay, intra-individual and inter-individual variation. Most of the intra-assay metabolite CV%s are less than 5%, indicating high robustness and accuracy of the urine NMR spectroscopy and the entire quantification process per se. The intra-individual metabolite variation over 30 days was large, with CV%s over 20% for the majority of metabolites and at the extreme 194% for sucrose and 225% for 2-furoylglycine. However, the population-based inter-individual metabolite variation was even larger, with CV%s over 40% for the majority of metabolites and at the extreme 585% for formate and 1655% for glucose (reflecting a positively skewed distribution of urinary glucose, partly due to several individuals with prediabetes and diabetes in NFBC66; **Figure 2**). These results indicate a sound base for epidemiological and genetic studies.

### Metabolite quantification in urine samples

Quantification of the 43 metabolites in the urine NMR spectra for the 1,004 people from the NFBC66 was done via semi-automated lineshape fitting analyses. This work is in progress and it will eventually be possible to provide quantifications for many additional metabolites; a preliminary list of over 100 assigned metabolites that we have identified is provided in **Supplementary Table S3**. However, this type of quantification approach is not feasible for routine applications at large-scale. Instead, an automated approach similar to the one we developed and adapted for quantifying serum and plasma lipid, lipoprotein and other metabolic information^8,16,18^ needs to be developed for the urine spectra. Proof of concept is shown with the initial results for fully automated quantitative regression analyses for creatinine and glucose (**Supplementary Figure S1** and **Figure 2**). As already known from serum NMR metabolomics, this type of approach for automated metabolite quantification works well.^8,9,23^

Therefore, it is likely that it will be possible to establish an optimised automated quantification model for most of the abundant urine metabolites. Regression-based spectral quantification methods are generally known to work well for heavily overlapping signal structures as typical for urine NMR spectra.^16,23^ With the regression modelling, quantification of all the metabolites in the urine NMR spectrum can be fully automated to take only a few seconds. Once the spectral data have been acquired for a sample, new identified and quantifiable metabolites can be retrospectively analysed (provided that sample preparations and experimental NMR settings are kept consistent). The final number of urine metabolites that may eventually be included is uncertain and will depend on multiple factors. In the current experimental set-up, we estimate it will be possible to automate the quantification for clearly more than 50 but likely not for all the metabolites listed in **Supplementary Table S3**.

It is estimated that there are around 3,000 compounds in the entire urine metabolome, of which there are currently 380 unique urine metabolites or metabolite species for which quantitative data are available.^1^ The approach described here to identify and quantify around hundred urine metabolites may therefore seem somewhat restricted. However, the quantitative serum NMR metabolomics platform is also limited to quantify “only” some 200 metabolic measures of the approximate 4,000 known serum compounds and this has not prevented novel metabolic measures being available for epidemiological and genetic studies with a plethora of new findings over the last few years.^9^ We anticipate that the quantitative urine metabolite data would lead to commensurate novelty in systems epidemiology. In epidemiology in particular it may be preferable to have a reasonable number of traits for as many people as possible, not vice versa.

### Preliminary epidemiological and genetic analyses

The lineshape fitting analyses of the urine NMR metabolomics data for the 1,004 individuals from NFBC66 allowed us to perform the first quantitative epidemiological analyses (**Table 1**). We also present some comparison to the serum NMR metabolomics data available for the same individuals (n=995). Some fundamental issues and corollaries are presented below.

### Metabolic associations

The correlations between the metabolites in urine (**Figure 4**) are generally weaker than the associations between most of the metabolic measures in serum (**Figure 3**). On average, the median of the absolute correlations between metabolites in urine was 0.10 (interquartile range, 0.04 – 0.19), and in serum 0.21 (0.08 – 0.44). These association characteristics are likely to partly reflect the larger intra-individual variation in the urine metabolites than those in serum, but they are also likely a sign of fundamental metabolic differences regarding serum and urine. The metabolic measures quantified via serum NMR metabolomics represent key systemic metabolic pathways (e.g., lipoprotein lipid metabolism) that are inherently physiologically correlated; it would not be expect to see drastic differences between individuals in these highly conserved biochemically essential metabolic pathways. However, in urine, which is a waste product, this type of tight inherent metabolic control is not necessary. Importantly, this might allow more specific biomarker findings from the urine data than would be possible from serum. Associations illustrated in **Figure 4** provide a proof of concept of the relevance of these novel quantitative urinary data. For metabolites in urine, positive correlations among amino acids, glycolysis- and citrate cycle-related metabolites, 3-hydroxyisobutyrate and 3-hydroxyisovalerate result in clear association clusters. The concentrations of the 8 amino acids are rather strongly correlated in urine with median correlation of 0.53 (interquartile 0.41 – 0.59). This is expected for these mostly apparently healthy individuals since, in healthy kidneys, the glomeruli filter all amino acids out of the blood and the renal tubules then reabsorb them back into the blood.

Metabolic correlations between urine and serum (**Figure 5**), with median absolute correlation of 0.04 (interquartile range, 0.02 – 0.07), are clearly weaker that the associations in serum (**Figure 3**) and in urine (**Figure 4**). However, several clearly detectable metabolic associations are present as elaborated in the caption for **Figure 5**. There is a clear excess of metabolic information by the combination of quantitative urine and serum metabolomics, illustrating an abundance of epidemiological novelty from quantitative urine metabolomics.

### Adiposity and urine metabolites

Associations between the 43 quantified urine metabolites (and the corresponding ones available in serum via the serum NMR metabolomics platform) and body-mass index (BMI) are illustrated in **Figure 6**. Despite rather large biological variation in the urine metabolite data, multiple associations are notable between the urine metabolites and BMI. While we ought to be cautious in interpreting cross-sectional associations, in comparison to recent work by Elliott and co-workers^4^ on urinary metabolic signatures of adiposity in two independent cohorts, the US and UK INTERMAP studies, we note multiple concordant associations for BMI, for example, negative with urinary p-Cresol sulphate and hippurate, and positive with 2-hydroxyisobutyrate and branched-chain amino acids isoleucine and valine, and aromatic amino acids tryptophan and tyrosine. The comparison of the amino acid results (e.g., valine and isoleucine) in urine and serum is of interest due to recent findings regarding the interplay between branched-chain amino acids, obesity, insulin resistance and the development of type 2 diabetes.^25-28^ For all the amino acids that are quantified from both urine and serum, the association direction with BMI is the same in serum and in urine; the association strengths however tend to be weaker in urine. As far as we are aware, these are the first results available combining quantitative metabolomics data from serum and urine at an epidemiological scale. We have previously illustrated, via Mendelian randomization analyses,^29^ that BMI is causally modifying circulating metabolism, including branched-chain amino acids.^30^ Potential causal effects of obesity on specific urine metabolites (e.g., via influences on kidney function) are largely unknown and will be one of our future aims of research with larger numbers of individuals. Even though urine is waste, urinary metabolites may serve as useful biomarkers (independent or together with serum metabolic measures) reflecting (patho)physiological effects of, e.g., obesity on systemic metabolism and organ function.

### Genome-wide analyses of urine metabolites

We performed a GWAS for the 43 quantified urine metabolites (**Table 1**). Despite having only 578 individuals available with both genome-wide data and quantified urine metabolites, we were able to replicate, at GWAS significance, two loci previously associated with the same urine metabolites.^31,32^ We found confirmatory evidence for formate, with chromosome 8 SNP rs4921913, associating at P= 4.59*10^−11^. The tagged region harbours arylamine N-acetyltransferase (*NAT2*) that is the candidate gene for this association. In addition, we confirmed an association of SNP rs1168674 in chromosome 12 with 2-hydroxyisobutyrate (P= 1.51*10^−22^). This region is in near vicinity to 4-hydroxyphenylpyruvate dioxygenase (*HPD*) that is likely involved in the metabolic pathway of 2-hydroxyisobutyrate.^31^ Manhattan plots for these associations are presented in **Figure 7**. These independently replicated genome-wide associations, with a very small number of individuals, are reassuring regarding the analytical processes of the presented urine NMR metabolomics platform.

### Statistical issues in epidemiology and genetics

Hundreds of metabolites have been identified in human urine samples with a combination of multiple spectroscopic technologies.^1,2,33,34^ However, most metabolomics applications have focused on dietary and various (biologically rather inapplicable) diagnostic issues typically with small numbers of individuals and profiling based analysis approaches.^35-37^ Comprehensive quantitative data on urine metabolites are rare,^2,38^ and if available, typically originate from other methods than NMR.^4,33,39^ This situation suggests that though the potential of urine NMR metabolomics is clearly recognised, the methodologies are still far from real-world large-scale applications. Nevertheless, we fully agree with Emwas and co-workers^2^ that molecular identification and absolute quantification are crucial both in epidemiology and genetics as well as if aiming to translate the biomarker discoveries to clinical practice.^8,38,40^ Therefore, a key characteristic for a urine NMR metabolomics pipeline will be that all metabolites are quantified in absolute terms. This means that in statistical analyses any platform output can be treated as any other clinical chemistry measure (e.g., glucose or cholesterol) in association testing and prediction models, including adjustments for appropriate confounding factors.^9^ The quantitative nature of the metabolite data makes this straightforward and also allows replication and meta-analyses across multiple studies.^40-42^ Here the joint analyses of urine and serum metabolomics data are a demonstration of how informative inherently simple quantitative molecular data can be. Notably, use of only spectral data would not enable the abovementioned molecular association analyses to be performed.

A general concern with using urine samples is metabolites that are not present in every sample or individual. This can actually be a high proportion of potentially detectable metabolites; from the automated quantitative analysis point of view this is a challenge calling for specific signal detection options. However, from the epidemiology point of view, it can be seen as a great opportunity. For example, specific drug-related metabolites may offer valuable epidemiological information as well as a base for potential pharmaceutical applications. Metabolites that would associate with certain foods or lifestyle factors, like smoking, would allow advantageous epidemiological approaches to be taken. Such metabolites may also indicate particular disease processes and could thereby provide specific clinically relevant biomarkers for risk assessment and early diagnoses. Here, only 4 out of the 43 quantified urine metabolites were absent for more than 10% of the samples, namely Sumiki’s acid, 2-furoylglycine, 3-(3-hydroxyphenyl)-3-hydroxypropanoate, and sucrose.

In addition to molecular quantification, systems epidemiology applications call for large numbers of individuals, at a minimum this is thousands, if not tens of thousands of individuals.^6,9^ The epidemiological data set described here for urine (43 metabolites quantified for 1,004 people) is already one of the largest in the area of quantitative urine metabolomics. These data, together with the various analytical tests, illustrate the key methodological and statistical characteristics of urine metabolomics. At the same time, however, this underscores that this field, particularly from the epidemiological perspective, is in its infancy. Therefore, the results presented here provide a good incentive to an open-access quantitative urine NMR metabolomics pipeline.

## Conclusions

Our quantitative analytical experimentation indicates high robustness and accuracy of the urine NMR spectroscopy methodology per se. The extensive epidemiological data illustrate clear inherent differences in the intra-fluid metabolic associations based on physiological and metabolic functions: the urine metabolites are in general only weakly interrelated, in contradistinction to highly correlated metabolic pathways represented by the quantitative serum data. The metabolic associations between serum and urine are weak, suggesting combining serum and urine metabolomics would increase the amount of independent metabolic information. While the intra-individual variation in urine metabolites is high, the even higher population-based inter-individual variation does provide a sound base for epidemiological and genetic applications. However, appropriate large-scale studies and replication data are crucial to enable statistically robust findings of biological relevance. The known genome-wide associations detected here with a very small number of individuals are reassuring for both the analytical process of the presented urine NMR metabolomics set-up and the intriguing potential of quantitative urine metabolite data in systems epidemiology. We anticipate this quantitative methodology to eventually offer a multitude of unique opportunities to study the role of urine metabolites, for example, in cardiometabolic health and diseases and as potential markers of kidney function. To the best of our knowledge, this project is novel both in the open access aspects and in the integrated large-scale systems epidemiology perspective that are likely to result in important epidemiological findings with high translational potential.

## Acknowledgements

QW was supported by a Novo Nordisk Foundation Postdoctoral Fellowship (grant number NNF17OC0027034). MG, SR, GDS and MAK work in a Unit that is supported by the University of Bristol and UK Medical Research Council (MC_UU_12013/1). JK was supported through funds from the Academy of Finland (grant numbers 297338 and 307247) and Novo Nordisk Foundation (grant number NNF17OC0026062). MHi was supported by the UK Medical Research Council Population Health Research Unit. MAK was supported by the Sigrid Juselius Foundation, Finland.

