## Supplementary Materials for "Proof of concept for quantitative urine NMR metabolomics pipeline for large-scale epidemiology and genetics"

### Data Supplement

**Northern Finland Birth Cohort of 1966**

**Figure S1.** The automated quantification of urinary creatinine and glucose from the NMR spectra.

**Table S1.** The automated sample preparation protocol for urine samples.

**Table S2.** Proton NMR spectroscopy measurement parameters and protocol for quantitative urine data.

**Table S3.** A preliminary list of assigned metabolites in urine NMR data.

**Northern Finland Birth Cohort of 1966**

The Northern Finland Birth Cohort of 1966 (NFBC66) included 12,058 children born alive into the cohort, comprising 96% of all births during 1966 in the region. Data collection in 2012 included clinical examination as well as serum and urine sampling at the age of 46 years for 5,788 and 4,549 individuals, respectively. The serum and urine samples were taken after overnight fasting. Informed written consent was obtained from all participants. The research protocols were approved by the Ethics Committee of University of Oulu and the Ethics Committee of Northern Ostrobotnia Hospital District, Finland. More information on the cohort and the 2012 data collection can be found at http://www.oulu.fi/nfbc.

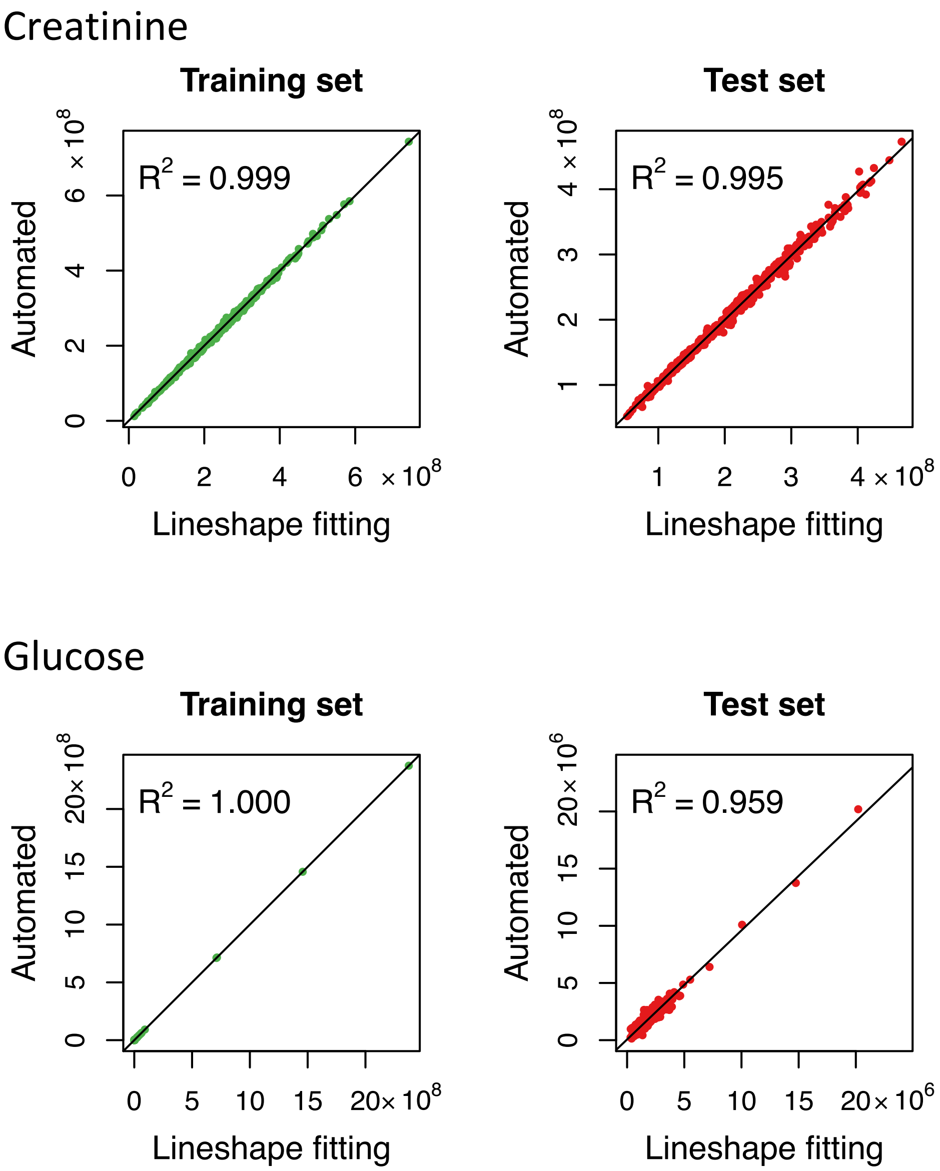


**Figure S1. The set-up and performance of the automated quantification of urinary creatinine and glucose from the NMR spectra.** The training set (n=500) illustrates how well the automated spectral regression analysis is reproducing the lineshape fitting analyses results for the absolute signal areas for both creatinine and glucose. The spectra in the test set (n=499) were analysed with the automated quantification models built with the training set. Comparisons to the lineshape fitting analyses results in the test set show excellent performance of the automated regression models for both metabolites. Ten lowest and 10 highest signal areas were included in the training set for both creatinine and glucose; the rest of the spectra for the training and test sets were chosen randomly. Multiple random modelling rounds gave essentially the same results indicating that the regression models are robust and very well defined.

**Table S1.** The automated sample preparation protocol for urine samples.

1. Overnight thawing of the samples in a refrigerator (+4°C).
2. Gentle mixing and centrifugation (3500 x g, 5 min, +4°C) of the thawed samples.
3. Transfer of 70 µl of phosphate buffer (1.5 M potassium dihydrogen phosphate, 0.2% sodium azide, 5.8 mM TSP in deuterium oxide, pH 7.0) to a 96 deep well plate by an automated liquid handler (PerkinElmer JANUS 8-tip Automated Workstation).
4. Transfer of 2 x 315 µl of the centrifuged urine samples to the deep well plate and mixing by aspirating and dispensing the sample (3 x 450 µl by JANUS).
5. Centrifugation (3500 x g, 5 min, +4°C) of the samples on the 96 deep well plate.
6. Transfer of 520 µl of the centrifuged samples to 5 mm 96 NMR tubes by JANUS.

Two control samples are prepared for each 96 NMR tube rack: Control sample "A" that is a real human urine sample (collected either from one person or pooled from samples taken from a few individuals) and a control sample "B" that is a simplified artificial urine sample containing only a simple mixture of a few urine metabolites and preservative (3.33 mM glucose, 10 mM creatinine, 0.5 mM alanine and 0.62 mM sodium azide in phosphate buffered saline). It is recommended to prepare, e.g., 100 ml of stock (10X) Control B solution (33.33 mM glucose, 100 mM creatinine, 5 mM alanine and 6.2 mM sodium azide in phosphate buffered saline) and dilute the solution with phosphate buffered saline when needed. A large number of 800 µl aliquots of both control samples are stored at -80°C. These quality control samples allow monitoring the quality and stability of sample preparations as well as NMR measurements. Automated quality control routines can be incorporated as part of the automated metabolite quantification software.

In the development of the sample preparation protocol, the inter-sample chemical shift variations arising from differences in sample conditions should be minimised. Previous efforts to solve this problem include sample acidification,[1] addition of phosphate buffer,[2] combined additions of potassium fluoride, phosphate buffer and deuterated ethylenediaminetetraacetic acid tripotassium salt ($K_3$EDTA-d12).[3] A high-throughput methodology calls for simplicity which may be achieved with the addition of reasonably strong phosphate buffer. Addition of potassium fluoride may cause only minor improvement but introduces the risk of breaking down some metabolites by addition of an acid. Addition of EDTA would introduce additional spectral signals that would complicate and even prevent some metabolite quantifications; in addition, some of the chemical shift variation would still remain.

**Table S2.** Proton NMR spectroscopy measurement parameters and protocol for quantitative urine data. These parameters have been optimized for a Bruker 600 MHz magnet equipped with a cryoprobe (Bruker Prodigy TCI 600 S3 H&F-C/N-D-05 Z).

| | |
|---|---|
| Pulse program | noesygppr1d |
| Size of fid | 81920 |
| Number of dummy scans | 0 |
| Number of scans | 16 |
| Sweep width | 21.0297 ppm |
| Acquisition time | 3.2440 s |
| Relaxation delay | 2.5 s |
| Receiver gain | 57 |
| Dwell time | 39.600 usec |
| Line broadening | 0.3 Hz |
| Temperature[a] | 295 K |

[a]Temperature calibration should be performed regularly (e.g. every 3 months) to ensure that the temperature settings are correct and all the samples are run at the same temperature. A 99.8% deuterated methanol sample in a sealed 5 mm NMR tube is commonly used to calibrate the temperature. A standard $^{1}H$ NMR spectrum is acquired and the fid is processed with a line broadening of 3.0 Hz. The probe temperature is calculated by measuring the distance between the $CH_3$ and OH signals (this is automatically performed using command “calctemp” in Topspin).

The first sample in the NMR tube rack is control sample A that is used to optimize the shimming routine. The samples are measured using Icon NMR automation and the SampleJet mode is 5 mm shuttle. The SampleJet sample changer unit is set to refrigerator temperature (+6°C) to enhance urine sample stability. Each sample has to wait 45 seconds in the magnet prior to operation to equilibrate the temperature. Five subsequent samples are automatically transferred to the SampleJet heater (set at 295.1 K) to pre-warm the samples close to the measurement temperature; this minimises the required equilibrating time in the magnet and speeds up the protocol. Once the temperature of the sample is stable, the signal is locked to the solvent, tuned and matched, as well as shimmed using the automated routine. Due to the high ion concentration of urine samples, a new solvent called “urine” was created in Topspin and several lock and loop parameters were optimised (Table S3). For comparison, the corresponding values for solvent $H_2O$ + $D_2O$ are shown below:

| Solvent | Lock power | Lock power instep | Loop gain | Loop time | Loop filter | Lock phase | Shift (ppm) |
|---|---|---|---|---|---|---|---|
| $H_2O + D_2O$ | -18 | 10 | -9.4 | 0.464 | 50 | 52.9 | 4.7 |
| Urine | -18 | 10 | -12.26 | 0.53 | 35 | 93 | 4.7 |

The noesygppr1d data (with the parameters below) are then acquired and processed using the automated routine. The NMR experimentation for one 96 NMR tube rack takes approximately 10 hours allowing more than 200 urine samples to be analysed in 24 hours.

Using this approach, deliberate decisions and compromises have been made to optimise for high-throughput and cost-effectiveness. The strategy is similar to that employed with the high-throughput quantitative serum NMR metabolomics platform.[4-6] The urine protocol uses 16 scans to acquire the NMR signal from the samples and applies a rather short relaxation delay of 2.5s. The latter affects only the smallest metabolites (e.g., formate and acetate) and if absolute concentration values (e.g., in relative to creatinine) are required, a small set of samples can be run with a longer, fully quantitative relaxation delay and the difference calibrated. If a cryoprobe is not available and/or the urine samples are particularly dilute, the number of scans can be temporarily increased. Within these experimental settings, a typical detection limit is around 1 µmol/l (or 0.2 µM/mM creatinine), depending on the signal multiplicity and amount of protons that yield the quantified signal.

**Table S3.** A preliminary list of assigned metabolites in urine NMR data.

| Metabolite |
| --- |
| 1-Methylhistidine |
| 1,3-Dimethylurate |
| 2-Furoylglycine |
| 2-Hydroxyisobutyrate |
| 2-Methyl-3-hydroxybutyrate |
| 2-Methylerythritol |
| 3-(3-Hydroxyphenyl)-3-hydroxypropanoate |
| 3-Aminoisobutyrate |
| 3-Hydroxybutyrate |
| 3-Hydroxyhippurate |
| 3-Hydroxyisobutyrate |
| 3-Hydroxyisovalerate |
| 3-Methylhistidine |
| 3-Methylxanthine |
| 4-Deoxythreonate |
| 4-Hydroxybenzoate |
| 4-Hydroxyhippurate |
| 4-Hydroxyphenylacetate |
| 4-Pyridoxate |
| 7-Methylxanthine |
| Acetate |
| Acetaminophen glucuronide |
| Acetaminophen sulphate |
| Acetoacetate |
| Acetone |
| Allantoin |
| Anserine |
| Arabinose |
| Ascorbate |
| Azelate |
| Beta-alanine |
| Betaine |
| Choline |
| Cis-aconitate |
| Citrate |
| Creatine |
| Creatinine |
| Dehydroascorbate |
| Dimethylamine |
| Dimethyl glycine |
| Dimethyl sulfone |
| Erythritol |
| Ethanol |
| Ethanolamine |

| |
|---|
| Formate |
| Fumarate |
| Glucose |
| Glucuronate |
| Glycine |
| Glycolate |
| Guanidoacetate |
| Hippurate |
| Hypoxanthine |
| Indoleacetate |
| Indoxyl sulphate |
| Isocitrate |
| Isopropanol |
| Isovalerylglycine |
| Kynurenate |
| L-Acetylcarnitine |
| L-Alanine |
| L-Asparagine |
| L-Carnitine |
| L-Fucose |
| L-Glutamate |
| L-Glutamine |
| L-Histidine |
| L-Isoleucine |
| L-Lactate |
| L-Leucine |
| L-Lysine |
| L-Methionine |
| L-Phenylalanine |
| L-Serine |
| L-Threonine |
| L-Tryptophan |
| L-Tyrosine |
| L-Valine |
| Mannitol |
| Methanol |
| Myoinositol |
| Methylamine |
| Methylmalonate |
| Methylsuccinate |
| N1-methyl-2-pyridone-5-carboxamide |
| N1-Methylnicotinamide |
| N-Acetylaspartate |
| N-Acetylneuraminate |
| N-Methylhydantoin |
| N,N-dimethylglycine |
| Orotate |

| |
| --- |
| Oxypurinol |
| p-Cresol sulphate |
| Pantothenate |
| Phenylacetylglutamine |
| Picolinate |
| Pimelate |
| Proline-betaine |
| Propionate |
| Propylene glycol |
| Pseudouridine |
| Pyrocatechol |
| Pyroglutamate |
| Pyruvate |
| Sarcosine |
| Scyllitol |
| Sebacate |
| Suberate |
| Succinate |
| Sucrose |
| Sumiki's acid |
| Tartrate |
| Taurine |
| Threonate |
| Trans-aconitate |
| Trigonelline |
| Trimethylamine |
| Trimethylamine N-oxide |
| Tyrosine |
| Uracil |
| Urea |
| Xylose |
